# Cytokine signaling converging on *IL11* in ILD fibroblasts provokes aberrant epithelial differentiation signatures

**DOI:** 10.1101/2022.12.20.521114

**Authors:** Miriam T. Kastlmeier, Erika Gonzalez Rodriguez, Phoebe Cabanis, Eva M. Guenther, Ann-Christine König, Lianyong Han, Stefanie M. Hauck, Fenja See, Sara Asgharpour, Christina Bukas, Gerald Burgstaller, Marie Piraud, Mareike Lehmann, Rudolf A. Hatz, Jürgen Behr, Tobias Stoeger, Anne Hilgendorff, Carola Voss

## Abstract

Interstitial lung disease (ILD) is a heterogenous group of lung disorders where destruction and incomplete regeneration of the lung parenchyma often results in persistent architectural distortion of the pulmonary scaffold. Continuous mesenchyme-centered, disease-relevant signaling likely initiates and perpetuates the fibrotic remodeling process, specifically targeting the epithelial cell compartment, thereby destroying the gas exchange area.

With the aim of identifying functionally relevant mediators of the lung mesenchymal-epithelial crosstalk that hold potential as new targets for therapeutic strategies, we developed a 3D organoid co-culture model based on human induced pluripotent stem cell-derived alveolar epithelial type 2 cells that form alveolar organoids in presence of lung fibroblasts from ILD patients as well as a control cell line (IMR-90). While organoid formation capacity and size was comparable in the presence of ILD or control lung fibroblasts, metabolic activity was significantly increased in ILD co-cultures. Alveolar organoids cultured with ILD fibroblasts further demonstrated reduced stem cell function as reflected by reduced *Surfactant Protein C* gene expression together with an aberrant basaloid-prone differentiation program indicated by elevated *Cadherin 2, Bone Morphogenic Protein 4* and *Vimentin* transcription.

In order to screen for key mediators of the misguided mesenchymal-to-epithelial crosstalk with a focus on disease-relevant inflammatory processes, we used mass spectrometry and characterized the secretome of end stage ILD lung fibroblasts in comparison to non-chronic lung disease (CLD) patient fibroblasts. Out of the over 2000 proteins detected by this experimental approach, 47 proteins were differentially abundant comparing ILD and non-CLD fibroblast secretome.

The ILD secretome profile was dominated by chemokines, including *CXCL1, CXCL3*, and *CXCL8*, interfering with growth factor signaling orchestrated by *Interleukin 11 (IL11)*, steering fibrogenic cell-cell communication, and proteins regulating extracellular matrix remodeling including epithelial-to-mesenchymal transition. When in turn treating alveolar organoids with *IL11*, we recapitulated the co-culture results obtained with primary ILD fibroblasts including changes in metabolic activity.

In summary, we identified mediators likely contributing to the disease-perpetuating mesenchymal-to-epithelial crosstalk in ILD. In our alveolar organoid co-cultures, we were able to highlight the importance of fibroblast-initiated aberrant epithelial differentiation and confirmed *IL11* as a key player in ILD pathogenesis by unbiased ILD fibroblast secretome analysis.

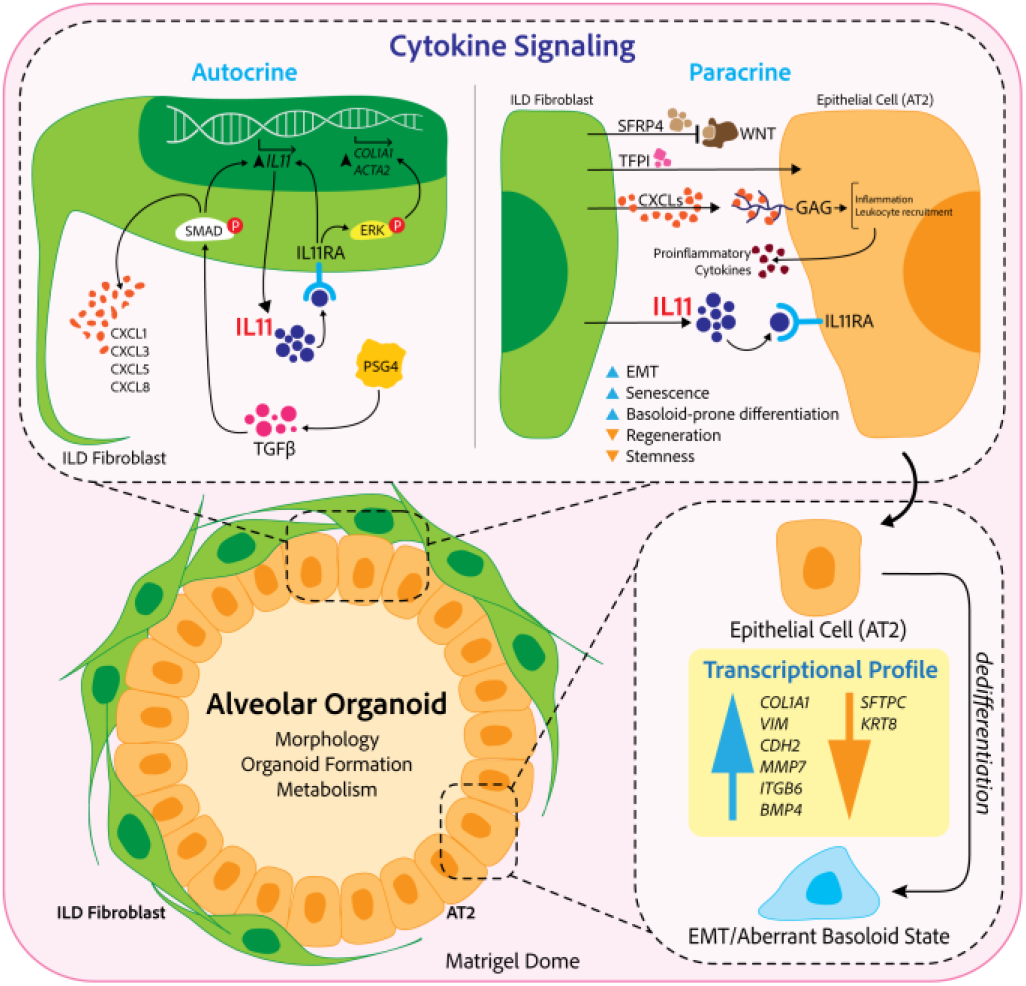

## 1 Introduction

Interstitial lung diseases (ILDs) comprise a variety of chronic pulmonary conditions that are characterized by structural remodeling of the gas exchange area (Glasser et al., 2010). ILD pathophysiology is centered on sustained inflammation and progressive scarring, ultimately resulting in irreversible tissue destruction and organ failure. Despite the exact pathogenesis of ILD still being unclear, genetic predisposition, age, sex and environmental exposure are known drivers of the disease (Lederer & Martinez, 2018; Strikoudis et al., 2019).

In ILD pathogenesis, fibroblast activation occurs through the impact of exogenous stimuli side-by-side with their activation through innate immune cells, especially monocytes and neutrophils, communicating via growth factor signaling and cytokine secretion. Subsequently, lactic acid release from fibroblasts as well as epithelial cells, induced by matrix metalloproteinases (MMPs), fibroblast growth factors and metabolic changes in turn further enhances fibroblast activation and accumulation (Dwyer et al., 2016). As a result, fibroblasts become the main loci of extracellular matrix (ECM) production and deposition. Induced by repeated inflammatory epithelial injury that leads to further leucocyte attraction and infiltration of the airspace and further perpetuating the pro-fibrotic circle of events, activated fibroblasts are discussed to induce epithelial-to-mesenchymal transition (EMT) in alveolar epithelial cells *via* SMAD and MAPK signaling (Kim et al., 2018), (O’Dwyer et al., 2016).

Driving and upholding the outlined pathophysiologic processes in ILD that ultimately result in severe tissue destruction and loss of the alveolar epithelium in end-stage ILD, a tightly knit crosstalk between the activated fibroblast and the damaged epithelium has been proposed (Ng et al., 2020; Ng et al., 2019; Yao et al., 2021). Here, the role of pathologic growth factor signaling and secreted cytokines such as *transforming growth factor ß (TGF-β)* and *Interleukin 17 (IL17)* were highlighted.

Adding to their detrimental role, activated fibroblasts – next to their capacity for pro-fibrotic signaling and EMT induction – have been shown to alter repair and regeneration of the injured gas-exchange area by affecting alveolar epithelial type 2 stem cell potential (Mou, 2021).

To address apparent knowledge gaps in the cytokine-driven, disease-relevant mesenchymal-to-epithelial crosstalk, we used lung organoids derived from human induced pluripotent stem cells (hiPSCs) through chemical directed differentiation into a sophisticated co-culture model. hiPSC-derived alveolar type 2 cells (iAT2s) are recognized as a useful tool to study lung diseases and regeneration capacities, and are recently emerging as a novel tool in environmental and occupational hazard assessment (Kastlmeier, Guenther, et al., 2022; Kong et al., 2021). iAT2-derived alveolar organoids recapitulate the characteristic three-dimensional (3D) structure of alveoli and are thereby ideal to mimic important functions of the gas exchange area *in vitro*. Their potential to study pulmonary disease *in vitro* (Hogan, 2021; Nikolić et al., 2018) is particularly versatile when targeting underlying molecular mechanisms (Strikoudis et al., 2019).

By the integration of this sophisticated methodology in a novel co-culture model, we were able to investigate the impact of primary ILD-derived lung fibroblasts on critical functions of hiPSC-derived alveolar organoids. The combination of this approach with unbiased secretome analysis allowed us to delineate functionally relevant signals of the pathologic crosstalk from the lung mesenchyme towards the alveolar epithelium with the aim to identify potential therapeutic targets.

## 2 Material and Methods

### 2.1 Human induced pluripotent stem cells (hiPSCs) and directed differentiation into lung progenitors

The hiPSC line BU3NGST was kindly provided by Prof. Darrell Kotton, Boston University, Center for Regenerative Medicine. This cell line is a dual-reporter construct composed of fluorochrome-encoding cassettes targeted to the endogenous NKX2.1 and SFTPC loci (BU3 NKX2.1^GFP^; SFTPC^tdTomato^) (Hawkins et al., 2017). hiPSCs were maintained in mTeSR1 (StemCell Technologies), on Matrigel (Corning) coated cell culture plates at 37 °C/5 % CO_2_ in a cell culture CO_2_ incubator. Cells were subcultured by using ReLeSR (StemCell Technologies) or Gentle Cell Dissociation Reagent (StemCell Technologies) (Jacob et al., 2017; Jacob et al., 2019).

BU3NGSTs were differentiated into NKX2.1^+^ lung progenitor cells and iAT2s as described previously by Jacob et al. (Jacob et al., 2017; Jacob et al., 2019). In short, hiPSCs were checked for their pluripotency *via* Alkaline Phosphatase staining (ES Cell Characterization Kit, CHEMICON International) or immunofluorescence staining of TRA 181 and SSEA 4 (ES cell characterization Kit, CHEMICON International). Induction of definitive endoderm was conducted *via* STEMdiff Definitive Endoderm Kit, (StemCell Technologies). On day 14 of differentiation, lung progenitor specification was evaluated by immunofluorescence staining of NKX2.1 (Invitrogen) and Albumin (ALB, R&D Systems). NKX2.1^GFP+^ lung progenitor cells were enriched by GFP signal for NKX2.1 based on a previously described protocol. The sorting was performed by FACS cell sorting at MACSQuant Tyto Cell Sorter (Miltenyi Biotec). For data evaluation FlowJo Version 7.2.1 and v10 was used. Purified lung progenitors were seeded in Matrigel (Corning) domes at a cell density of 50 cells/μL and passaged every second week. To increase SFTPC^tdTomato+^ cells CHIR withdrawal and addback was performed. At day 45 of differentiation, iAT2s were enriched by flow cytometry (MACSQuant Tyto Cell Sorter, Miltenyi Biotec) using tdTomato signal for SFTPC expression and subsequently cultured as 3D alveolar organoids. Differentiated SFTPC^tdTomato+^ iAT2s in 3D Matrigel were grown in CK+DCI medium, with media changes every 48 – 72 h. Alveolar organoids were passaged every 14 days.

### 2.2 Primary human fibroblast culture

Primary human lung fibroblasts from ILD patients and non-CLD controls (P4) for co-culture experiments (ILD fibroblasts) and MS based secretome analysis (ILD and non-CLD fibroblasts) were isolated according to a published protocol (Heinzelmann et al., 2018) and obtained through the CPC-M bioArchive at the Comprehensive Pneumology Center in Munich, Germany. All patients underwent surgery at the LMU Hospital and the Asklepios Pulmonary Hospital Munich-Gauting. Tissue from ILD patients (n = 3, Table 1) was provided through lung transplantation. Control fibroblasts were derived from lung tissue resections of age-matched non-CLD patients (female n = 1, = 3, male n = 2).

**Table 1.** Patient characteristics

| Patient | Diagnosis | Age at sample collection (years) | Sex | Surgery | Smoking history |
| --- | --- | --- | --- | --- | --- |
| 1 | Connective Tissue Disease related ILD | 48 | F | Lung transplantation (single) | no |
| 2 | Idiopathic pulmonary fibrosis | 62 | M | Lung transplantation (single) | no |
| 3 | Hypersensitivity Pneumonitis, Rheumatoid arthritis | 57 | F | Lung transplantation | no |

The study was approved by the local ethics committee of the Ludwig-Maximilians University of Munich, Germany (Ethic vote #333-10) and written informed consent form was obtained for all study participants.

Human fetal lung fibroblasts (IMR-90, P8) for control co-cultures were obtained from ATCC (Catalog # CCL-186™) and grown in Dulbecco’s Modified Eagle Medium: Nutrient F-12 (DMEM/F12; Gibco) with 20 % fetal bovine serum (FBS SUPERIOR, Sigma) and 1 % penicillin-streptomycin (Pen Strep, Gibco).

Cells were seeded in 6-well plates. at a density of 1×10^5^ cells in 2 mL media (DMEM/F12, 20 % FBS, 1 % penicillin/streptomycin) per well until reaching 80 % confluency. Once confluent, each well was washed three times with a 15-minute incubation per wash with 1 mL of FBS-free culturing medium (DMEM/F12, 1 % penicillin/streptomycin) to eliminate remaining FBS. Fibroblasts were cultured for 48 h in FBS-free medium, supernatants were collected and stored at -80 °C for further analysis.

### 2.3 Secretome analysis by mass spectrometry

#### Sample preparation for proteomics

Each 500 μL supernatant was subjected to tryptic digest applying a modified filter aided sample preparation procedure (Grosche et al., 2016; Wiśniewski et al., 2009). After protein reduction and alkylation using DTT and iodoacetamide, samples were denatured in UA buffer (8 M urea in 0.1 M Tris/HCl pH 8.5) and centrifuged on a 30 kDa cut-off filter device (PALL or Sartorius) and washed thrice with UA buffer and twice with 50 mM ammoniumbicarbonate (ABC). Proteins were proteolysed for 2 h at room temperature using 0.5 μg Lys-C (Wako) and subsequently for 16 h at 37 °C using 1 μg trypsin (Promega). Peptides were collected by centrifugation and acidified with 0.5 % trifluoroacetic acid.

#### Mass spectrometric measurements

LC-MSMS analysis was performed on a Q-Exactive HF mass spectrometer (Thermo Scientific) each online coupled to a nano-RSLC (Ultimate 3000 RSLC; Dionex). For subsequent analysis on the Q-Exactive HF, tryptic peptides were accumulated on a nano trap column (300 μm inner diameter × 5 mm, packed with Acclaim PepMap100 C18, 5 μm, 100 Å; LC Packings) and then separated by reversed phase chromatography (nanoEase MZ HSS T3 Column, 100 Å, 1.8 μm, 75 μm × 250 mm; Waters) in a 80 minutes non-linear gradient from 3 to 40 % acetonitrile in 0.1 % formic acid at a flow rate of 250 nL/min. Eluted peptides were analyzed by the Q-Exactive HF mass spectrometer equipped with a PepSep PSS1 source. Full scan MS spectra (from m/z 300 to 1500) and MSMS fragment spectra were acquired in the Orbitrap with a resolution of 60.000 or 15.000 respectively, with maximum injection times of 50 ms each. The up to ten most intense ions were selected for HCD fragmentation depending on signal intensity (TOP10 method). Target peptides already selected for MS/MS were dynamically excluded for 30 seconds. Data are available via ProteomeXchange with identifier PXD039554 (Deutsch et al., 2019; Perez-Riverol et al., 2021).

#### Protein Identification and label-free quantification

Proteome Discoverer 2.5 software (Thermo Fisher Scientific; version 2.5.0.400) was used for peptide and protein identification via a database search (Sequest HT search engine, SequestHT score:1) against Swissprot human database (Release 2020_02, 20432 sequences), considering full tryptic specificity, allowing for up to two missed tryptic cleavage sites, precursor mass tolerance 10 ppm, fragment mass tolerance 0.02 Da. Carbamidomethylation of Cys was set as a static modification. Dynamic modifications included deamidation of Asn, Gln and Arg, oxidation of Pro and Met; and a combination of Met loss with acetylation on protein N-terminus. Percolator was used for validating peptide spectrum matches and peptides, accepting only the top-scoring hit for each spectrum, and satisfying the cutoff values for false discovery rate (FDR) < 5 %, and posterior error probability < 0.01.

The quantification of proteins was based on abundance values for unique peptides. Abundance values were normalized on total peptide amount and protein abundances were calculated summing up the abundance values for admissible peptides. The final protein ratio was calculated using median abundance values. The statistical significance of the ratio change was ascertained employing the T-test approach described in Navarro et al., 2014 (Navarro et al., 2014), which is based on the presumption that we look for expression changes for proteins that are just a few in comparison to the number of total proteins being quantified. The quantification variability of the non-changing “background” proteins can be used to infer which proteins change their expression in a statistically significant manner. Proteins with increased or decreased abundance were filtered with the following criteria: proteins were considered to be decreased in abundance below an abundance of ratio of 0.5 fold and increased abundance above 2 fold, proteins identified with a single peptide were excluded and just significant proteins were considered (P value < 0.05, P values were adjusted for multiple testing by Benjamini-Hochberg correction). Additionally, at least two MSMS identifications had to be identified to include the protein ratio.

#### Enrichment analysis

Pathway enrichment analyses were performed in Cytoscape (3.9.0) with the ClueGo plugin (v2.5.8) for significantly increased or decreased proteins. The following ontologies were used: KEGG (8093), GO_MolecularFunction-EBI-UniProt (18336), GO_BiologicalProcess-EBI-UniProt (18058). Accession IDs were used as identifiers and the analysis was performed with the standard software settings provided in the ClueGo app (Bindea et al., 2009).

### 2.4 Mesenchymal-epithelial co-culture

Primary lung ILD fibroblasts and IMR-90 (control fibroblast cell line) were grown in cell culture flasks until 70 % confluency. A single cell suspension was prepared using 0.25 % EDTA-Trypsin (Gibco). iAT2s were grown into alveolar organoids for up to two weeks in Matrigel domes. Single cell suspension was obtained with Dispase (Corning) and 0.25 % EDTA-Trypsin as described by Jacob et al. (Jacob et al., 2019). Human ILD and IMR-90 fibroblasts as well as iAT2s were counted and directly seeded either in equal 1:1 (F_low_) or 1:5 (F_high_) iAT2s to fibroblasts seeding densities in undiluted Matrigel domes in 8-chamber wells (20 μL Drops, Falcon), 96-well plates (50 μL Drops, Greiner) or 12-well plates (50 μL Drops, Greiner). Co-cultures used CK+DCI media that was changed every 48 h to 72 h for up to 12 days of cultivation.

### 2.5 Immunofluorescence microscopy

3D alveolar organoids mono- and co-cultures as described in section 2.1 (mono-culture) and 2.4 (co-culture) were cultured in 8-chamber wells for immunofluorescence analysis (Nunc Lab-Tek Chamber Slide System, 8-well, Permanox slide, 0.8 cm^2^/well). After alveolar organoids were formed, fixation was achieved with ice cold methanol and acetone (1:1v/v) for 5 minutes at -20 °C. Cells were washed with PBS and stained with the respective primary antibody in buffer containing 0.1 % BSA and 0.1 % Triton X-100 overnight at 4 °C. The next day, cells were washed 3 times with PBS and incubated in buffer with the respective fluorescent conjugated secondary antibody at a dilution of 1:500 and DAPI diluted 1:1.000 overnight at 4 °C. The following day, cells were washed gently, growth camber removed and remaining microscope slide mounted with fluorescent mounting media (Dako) and covered with a coverslip. Slides were stored at 4 °C until imaging. Imaging was performed using a confocal laser scanning microscope (CLSM) Zeiss LSM 880 with Airyscan and edited afterwards using ZEN 2.5 software (Zeiss). Detailed information on the primary and secondary antibodies are given in Supplementary Table 1.

### 2.6 Operetta high content imaging and Napari organoid counter

Live imaging of all alveolar organoid mono- and co-cultures was performed using the Operetta CLS high-content analysis system (Operetta CLS, PerkinElmer) at time points 5, 8 and 12 days during the co-culture experimental set-up. Pictures were analyzed by the Harmony 3.5.2 high-content imaging and analysis software with PhenoLOGIC. Multi-plane confocal 3D images were visualized in Napari image viewer (Python) as maximum intensity projections and the automatic measurements obtained from the “Napari organoid counter” (Bukas, 2022) were visually checked and manually curated, resulting in output of size and numbers of formed alveolar organoids between iAT2s and human fibroblasts (ILD and control IMR-90). The Canny Edge Detection (Canny, 1986) algorithm is used for identifying the organoids, while pre- and post-processing steps have been included, to ensure the image matches the detector’s expected input and the number of detected organoids along with their size is returned. More specifically, the organoids are approximated to ellipses and the algorithm fits orthogonal bounding boxes around each, with the height and width of each box corresponding to the two diameters of the organoid which are then in turn used to approximate the object’s area.

### 2.7 Quantitative real-time PCR

Co-cultures were lysed in RLT Plus Lysis Buffer (Qiagen) and RNA isolation was performed with the RNeasy Mini Kit (Qiagen) according to the manufacturer’s instructions. Cell lysis from organoids and co-culture assays was performed with peqGOLD TriFast (VWR Life Science) as recommended by the manufactures followed by RNA isolation with the RNeasy Mini Kit (Qiagen). RNA was transcribed into cDNA by reverse transcriptase using the High-Capacity cDNA Reverse Transcription Kit (Thermo Fisher Scientific) according to the manufacturer’s instructions. 5 ng of cDNA was added to a final concentration volume of 10 μL, Random Nonamers (Metabion) and master mix (Invitrogen, Thermo Fisher Scientific) was added to each RNA sample. cDNA was diluted with ultrapure H_2_O. qPCR was performed in 96-well format using the quantitative real-time PCR System (Roche 480 LightCycler). 2 μL cDNA were added to a final reaction volume of 10 μL containing H_2_O, 480 SYBR Green (LightCycler, Roche Diagnostics) and the primer mix (100 μM). Gene expression was normalized to *ß-Actin* control for genes *Vimentin* (*VIM*), *Integrin Subunit Beta 6* (*ITGB6*) and *Cadherin 2* (*CDH2*), and normalized to an average of *ß-Actin* and *HRPT* control for genes *Surfactant Protein C* (*SFTPC*), *Keratin 8* (*KRT8*), *Collagen 1A1* (*Col1A1*), *Matrix Metallopeptidase* (*MMP7*) and *Bone Morphogenic Protein* 4 (*BMP4*), the fold change was calculated using the 2^ (-ddC) method. Sequence information of used primers are given in Supplementary Table 2. Data obtained from qPCR are presented relative to respective control co-cultures, to demonstrate influence of disease background.

### 2.8 Metabolic activity estimated by WST-1 assay

WST-1 assays were performed at day 2, 3, 5 and 7 of alveolar organoid co-cultures (human ILD or IMR-90 control fibroblasts). WST-1 reagent (Roche Diagnostics) was added to the culture medium in a 1:10 dilution. The culture medium was used as background control. After 2 h of incubation at 37 °C and 5 % CO_2_, 100 μL of media from every sample was transferred to a microplate (Thermo Scientific; Fisher Scientific) and the absorbance of the sample against the background was measured with a TECAN reader (TECAN; infinite M200 PRO).

## 3 Results

### 3.1 Fibroblast induced changes in organoid formation and metabolic activity in co-cultured alveolar organoids

An overview of the experimental workflow is provided in Supplementary Figure 1A-C. iAT2s (Figure 1A) were successfully co-cultured of with both ILD and IMR-90 control fibroblasts, resulting in the formation of proliferative alveolar organoids (Figure 1B). Cell-cell contact of iAT2s and fibroblasts in (ILD) co-culture was demonstrated by partial encapsulation of alveolar organoids by α-SMA expressing fibroblasts (Figure 1C, white arrows).

**Figure 1.**
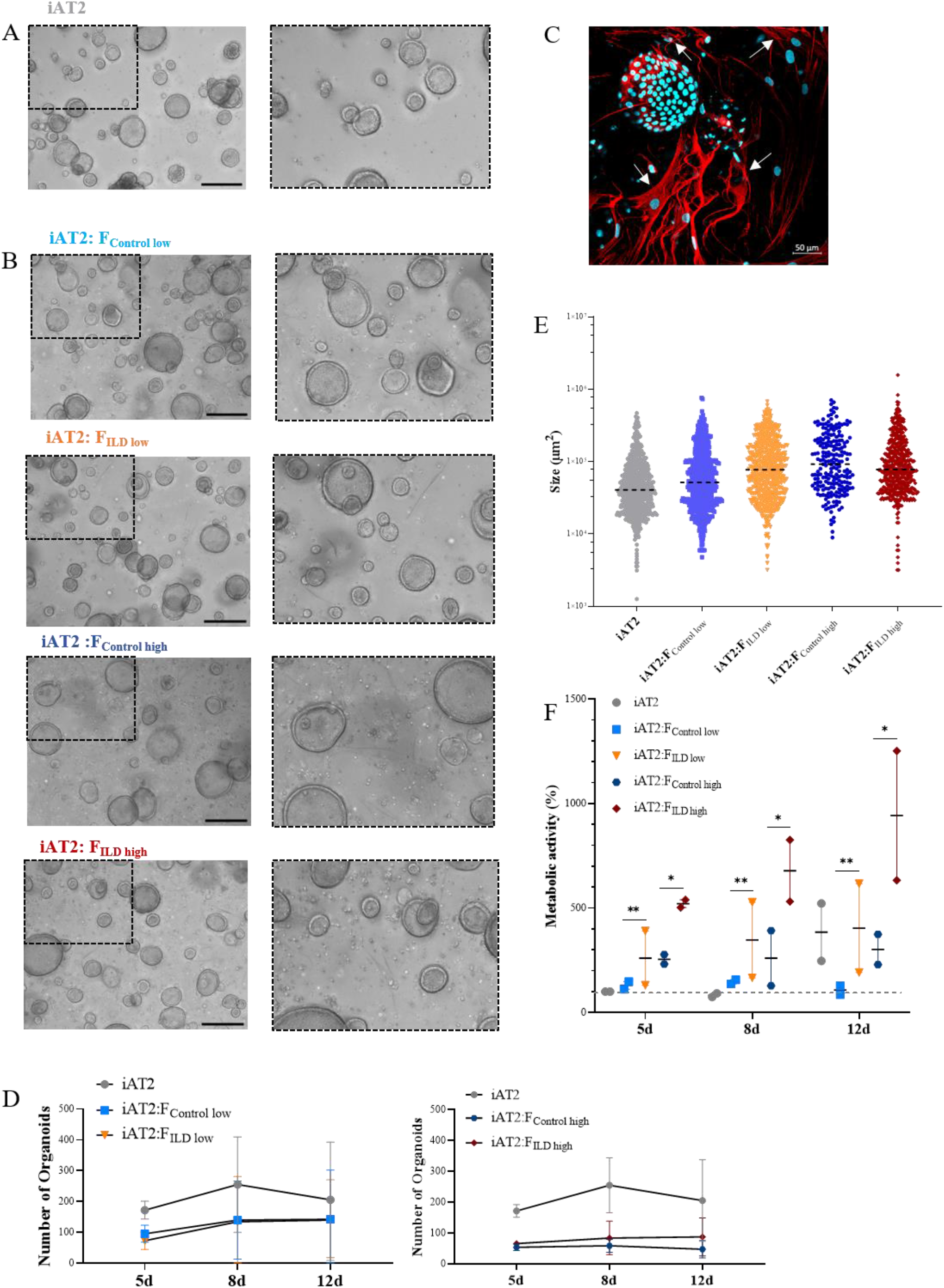
(A) Representative maximum intensity projections from high-content images of iAT2s growing as alveolar organoids at day 12. Scale bar 500μm. (B) Immunofluorescence of 3D co-culture of iAT2s with ILD fibroblasts at day 12 (α-SMA: red, DAPI: cyan). Scale bar 50 μm. (C) Representative maximum intensity projections from high-content images of different co-culture conditions showing iAT2s growing with human fibroblasts for 12 days. Scale bar 500 μm. Zoom-ins show a 3x optical magnification. (D) Scatter plots (dashed black lines: median) indicate size (μm2) of organoids in co-cultures at day 12 across three independent biological replicates. (E) Number of formed organoids in co-cultures at day 12, N = 3. (F) Metabolic activity of co-cultured organoids at 5, 8 and 12 days of co-culture. Results show the increase in percentage across two biological replicates in comparison to d5 iAT2 organoids alone, representing the baseline of 100% metabolic activity (dashed grey line across dataset). Statistics: unpaired t-Test, *p<0.05, **p<0.01.

Image analysis of the 3D co-cultures revealed a reduction in organoid formation capacity in the presence of human fibroblasts (Figure 1D). Although alveolar organoid size was not significantly affected by fibroblast co-culture (Figure 1B, E), quantitative assessment of organoid size (area, μm^2^) and number of images obtained from alveolar organoids and co-cultures with either ILD or IMR-90 control fibroblasts in 1:1 (F_ILD/Control low_) or 1:5 (F_ILD/Control high_) seeding density (Figure 1B, D) demonstrated a negative correlation of fibroblast seeding density (ILD or IMR-90) with the alveolar organoid formation capacity.

In contrast, co-culture with ILD fibroblasts significantly increased metabolic activity of alveolar organoid in comparison to IMR-90 control co-cultures (Figure 1F).

### 3.2 Presence of ILD fibroblasts leads to aberrant epithelial gene expression changes

In order to relate the observed changes in organoid formation capacity and metabolic activity (Figure 1) to changes in gene expression, we measured critical markers of stem cell function and epithelial differentiation in co-cultured organoids.

Indicating changes in (stem) cell function and epithelial injury, we showed decreased expression of the alveolar stem cell marker *SFTPC* in the presence of ILD fibroblasts in both seeding ratios (F_low_ and F_high_; Figure 2A). In line with this, *Keratin 8* (*KRT8*) expression levels were reduced under the impact of ILD primary fibroblasts (Figure 2A). Further, the distal epithelial marker *Integrin Subunit Beta 6* (*ITGB6*) as well as *Bone Morphogenetic Protein 4* (*BMP4*) showed increased transcription in ILD co-cultures when high seeding densities were applied (Figure 2A). Genes associated with regulation of extracellular matrix formation and remodeling including *Collagen 1A1* (*Col1A1*), *N-Cadherin 2* (*CDH2*) and *Vimentin* (*VIM*) showed increased expression in alveolar organoids co-cultured with ILD fibroblasts in high seeding ratios (Figure 2B). Likewise, expression levels of *Matrix Metallopeptidase 7* (*MMP7*) were increased in ILD co-cultures compared to control co-cultures. (Figure 2B).

**Figure 2.**
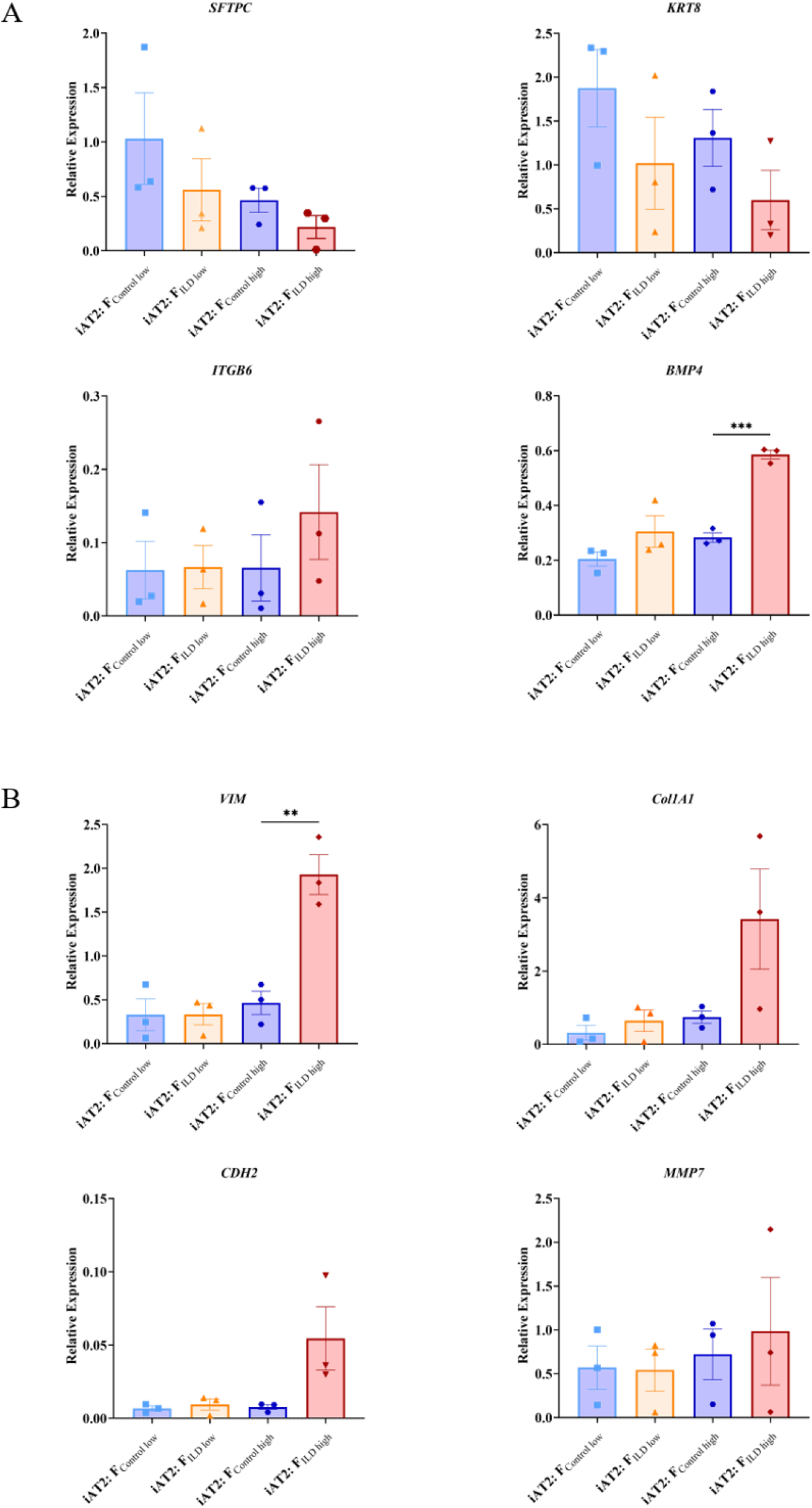
Relative gene expression at day 12 measured in AT2s co-cultures with ILD or IMR90 control fibroblasts in two seeding densities (high or low) compared to reference gene expression (HK; average of ß-Actin *(ACTB)* and hypoxanthine guanine phosphoribosyl transferase (*HRPT)*). (A) Epithelial and stem cell markers and (B) Genes associated with aberrant differentiation of epithelium. N = 3, unpaired t-Test, **p<0.005, ***p<0.0005

### 3.3 ILD fibroblast secretome reveals proinflammatory signaling converging on *IL11* stimulating epithelial remodeling

To characterize fibroblast driven communication resulting in gene expression and phenotypical changes in the alveolar epithelium in ILD, supernatants of ILD and non-CLD fibroblasts were subjected to mass spectrometry (MSMS).

MS analysis detected an overall of 2625 expressed proteins, of which 47 were significantly more and 55 significantly less abundant when comparing ILD-derived fibroblast to non-CLD controls (Supplementary Table 3, 4, Figure 3A). The top 15 differentially expressed proteins (increased and decreased abundance) are listed in Supplementary Table 1 and 2.

**Figure 3.**
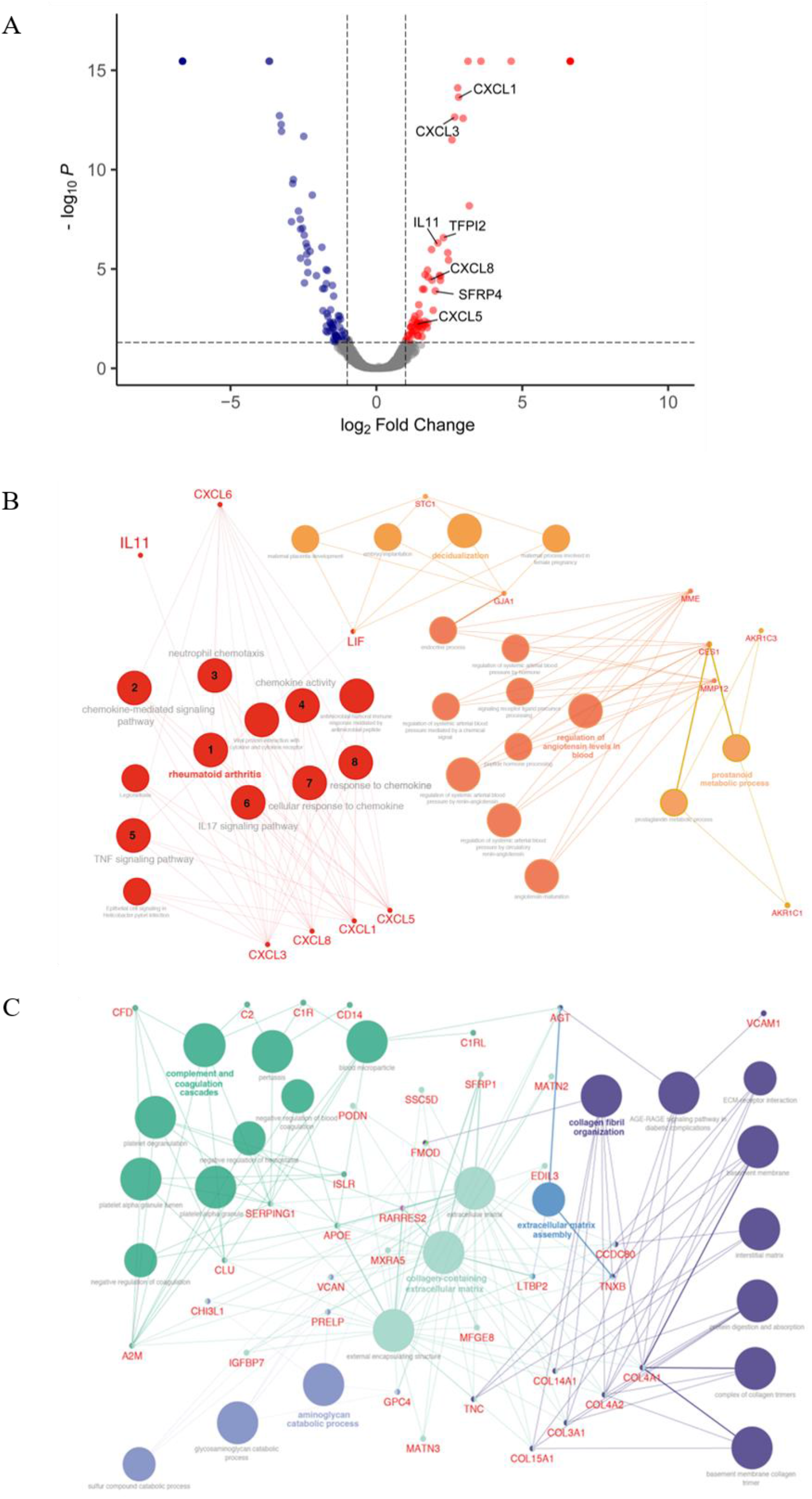
Differential protein expression comparing ILD fibroblasts (F_ILD_) vs. non-chronic lung disease fibroblasts (F_control_). (A) Volcano plot visualizing significantly regulated proteins (47 up, 55 down) detected by MS secretome analysis of ILD or non-CLD control fibroblasts. Data showing log_2_ fold change against the adjusted P value [log10]. Significantly upregulated proteins are depicted in red and significantly downregulated proteins in blue. (Total: 102 significantly regulated proteins with 5% FDR < 0.05, adj. p_value < 0.05). Pathway enrichment and protein interaction network of proteins with (B) increased and (C) decreased abundance using the Cytoscape plugin ClueGo. The following ontologies were used: KEGG, molecular functions and biological processes. The connectivity of the pathways is described by functional nodes and edges that are shared between proteins with a kappa score of 0.4. Only enriched pathways are visualized and the node size indicates the p-value (*p*-value ≤ 0.05). Proteins from the same pathway share the same node color and the bold fonts indicate the most important functional pathways that define the names of each group. Enriched Pathways: 1. rheumatoid arthritis, 2. chemokine-mediated signaling pathway, 3. neutrophil chemotaxis, 4. chemokine activity, 5. *TNF* signaling pathway, 6. *IL17* signaling pathway, 7. cellular response to chemokine, 8. response to chemokine.

Proteins with increased abundance predominantly belonged to the C-X-C motif chemokine family (*CXCL1, CXCL3*, and *CXCL8*), as well as to the interleukin family (*IL13RA, IL11*) and included gap junction proteins (*connexin 43, GJA1*). Further, *Pregnancy Specific Beta-1-Glycoprotein 4 (PSG4)* as well as WNT signaling modulator *SFRP4* were found amongst the top 15 proteins.

Accordingly, pathway enrichment analysis of proteins with increased abundance classified the responses as cytokine activity, chemokine-mediated signaling pathway and TNF-signaling pathway. Furthermore, cellular/response to chemokines, *IL17* signaling pathways and neutrophil chemotaxis were identified, indicative of a strong inflammatory response.

ClueGo, a Cytoscape plug-in for network analysis, highlighted proteins associated with rheumatoid arthritis, a disease often complicated by the development of lung fibrosis and characterized by the presence of inflammatory chemokines (Figure 3B).

Proteins with decreased abundance in the in ILD secretome were dominated by candidates involved in ECM production, ECM assembly or ECM reorganization as well as coordination of myofibroblast differentiation (*PDGFRL*) together with a downregulation of proteins involved in complement and coagulation cascade pathways (Figure 3C).

### 3.4 *IL11* acts as a driver for aberrant signatures in hiPSC-derived alveolospheres

Based on the MSMS secretome analysis, *IL11* emerged as a top player in the mesenchymal-to-epithelial disease crosstalk. In consequence, we exposed alveolar organoids (Figure 4A) to *IL11* in order to investigate its functional relevance.

**Figure 4.**
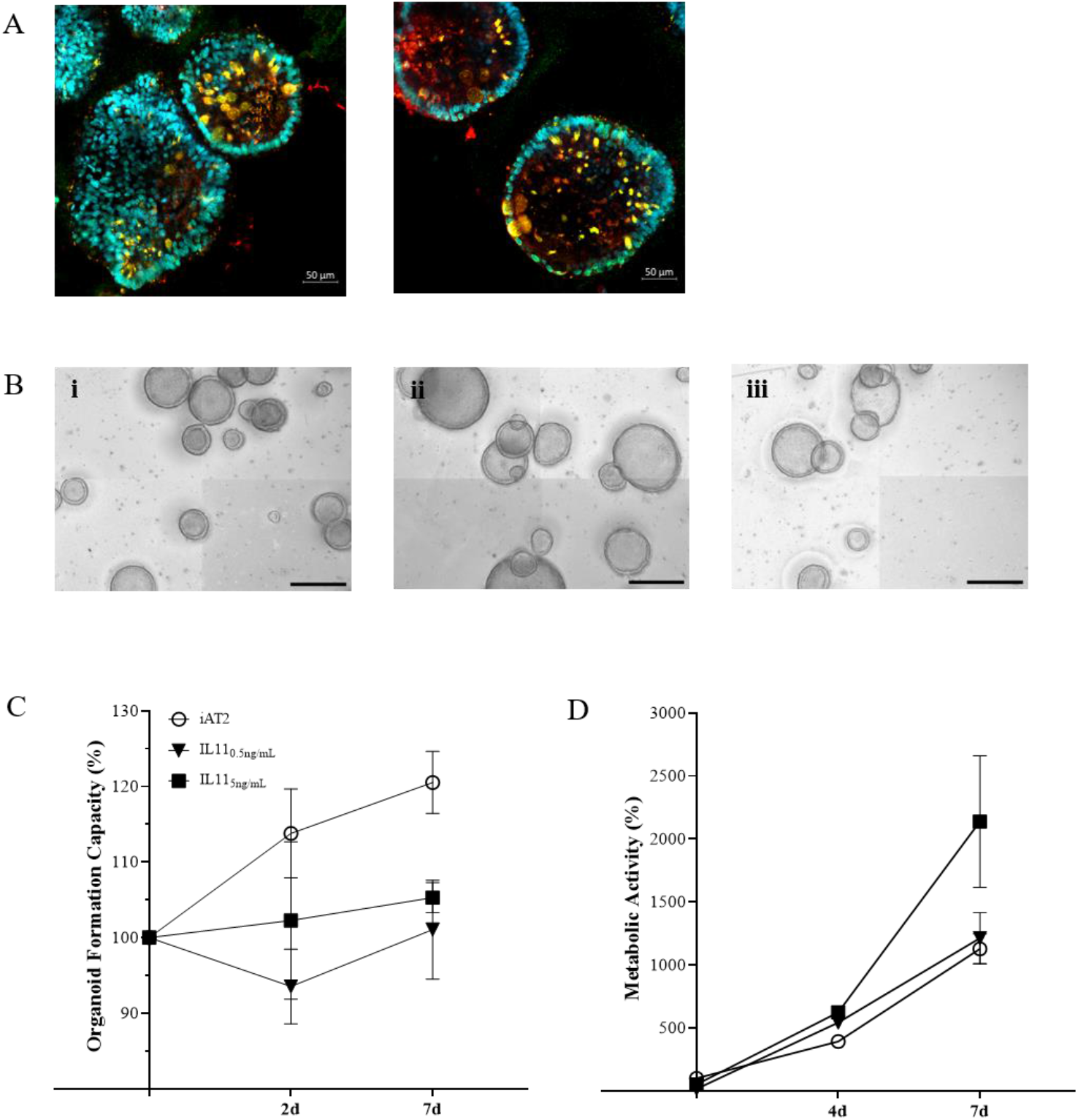
(A) Immunofluorescence of untreated alveolar organoids at day 14 of culture (*SFTPC*: red, *NKX2*.1: green, DAPI: cyan). (B) Representative maximum intensity projections from high-content images of (i) untreated alveolar organoids, (ii) 0.5 ng/mL or (iii) 5 ng/mL *IL11* treatment. Treatment started at day 7 of culture every 48h. Scale bar, 500 μm. (C) Organoid formation capacity of alveolar organoids treated with 0.5 or 5ng/mL *IL11*. (D) Metabolic activity of alveolar organoids treated with 0.5 or 5ng/mL *IL11*. (C&D) Each value is graphed as percentage of the respective starting culture at day 7 set to 100%.

Experiments (Dose_low_ = 0.5 ng/mL, Dose_high_ = 5 ng/mL) (Figure 4B - D) recapitulated the results observed in epithelial-fibroblasts co-cultures (section 3.1), *i*.*e*., we demonstrated reduced organoid formation capacity (Figure 4B (iii), 4C) and increased metabolic activity (WST-1) in *IL11* treated alveolar organoids (treatment from day 7 - 14 of culture) (Figure 4D). In addition, treatment of growing alveolar organoid monocultures with *IL11* (20 ng/mL) led to an increase in alveolar organoid size followed by apoptosis within 5 days of culture.

## 4 Discussion

In ILD, sustained inflammation and scarring of the gas exchange area ultimately result in destruction of the pulmonary scaffold and organ failure. Excessive deposition of ECM as well as epithelial damage and dedifferentiation is widely transmitted by the misguided interaction of fibroblasts and epithelial cells (Lewis et al., 2018). Therefore, improved understanding of the mesenchymal-to-epithelial crosstalk remains a centerpiece in finding new avenues to monitor and treat ILD. However, signaling factors with functional relevance and their distinct role in ILD pathogenesis remain understudied.

This study aimed at deciphering disease-relevant candidates of mesenchymal-to-epithelial crosstalk that could serve as potential targets for future therapeutic strategies. By advancing a sensitive human iPSC-derived alveolar organoid culture into a human fibroblast co-culture model, we successfully demonstrated the importance of fibroblast-driven, cytokine-centered signaling in inducing the impairment of key epithelial cell functions, including differentiation and metabolism. In combination with an unbiased proteomic approach, we were able to identify important mediators that translate these effects such as *IL11*, one of the top 15 proteins with increased abundance in ILD. The previously identified role of *IL11* in chronic inflammatory lung diseases in line with its potential to drive pro-fibrotic mesenchymal-epithelial crosstalk, supported the relevance of our approach on the one hand, while we were able to contribute the important functional consequences of its signaling in our human alveolar organoid model on the other (Cook & Schafer, 2020; Ng et al., 2020; Ng et al., 2019).

In our *in vitro* co-culture approach, primary human pulmonary fibroblasts and iAT2s formed alveolar organoids that successfully recapitulated tissue characteristics of the distal lung in three dimensions, in contrast to 2D, plastic culture conditions (Figure 1). Primary human fibroblasts, both ILD-derived and IMR-90 control cells, demonstrated effects that correlated with seeding density leading to reduced organoid number and increased organoid size after 12 days of co-culture. The findings indicate that the co-culture with fibroblasts per se is able to change the microenvironment of iAT2s, thereby impacting organoid formation. As studies indicated, that the activation of primary lung fibroblasts in a disease-comparable fashion is easily achieved (Kathiriya et al., 2022), our findings likely explain the disease-independent fibroblast effects in alveolar organoids.

In contrast, the significant increase in metabolic activity was provoked by diseased human fibroblasts in high-seeding ratios, indicating their potential to impact on the metabolic program of the epithelial cell, potentially indicating epithelial dedifferentiation or EMT (Kalluri, 2009; Kalluri & Weinberg, 2009; Wang et al., 2020). These considerations are supported by the expression signature characterizing the co-cultures: The decrease in *SFTPC* expression (Adams et al., 2020; Katzen & Beers, 2020; Kortekaas et al., 2022; Kortekaas et al., 2021) we observed in the ILD co-cultures points towards the loss of epithelial stem cell characteristics, as *SFTPC* expression is sensitive to epithelial inflammation and injury, lately supported by studies in Sars-CoV-2 infected alveolar organoids (Mou, 2021). Differentiation of AT2s is key for regeneration in injured alveoli marked by their expression of transient basaloid features such as *Keratin 5* (*KRT5*) and the amount of alveolar *KRT5*^*+*^ basaloid cells directly correlates with disease progression in pulmonary fibrosis (Kathiriya et al., 2022; Khan et al., 2022; Ng et al., 2022), mirrored by the decreased expression observed in our study. Closely related, downregulation of *KRT8*, an important marker of AT2 to (pre)AT1 transdifferentiation during epithelial regeneration (Ng et al., 2022; Strunz et al., 2020) was associated with the increased expression of EMT markers in a murine bleomycin lung injury model including pro-fibrogenic proteins such as *ITGB6* (Strunz et al., 2020). In line with these findings, we demonstrated increased *ITGB6* expression in ILD co-cultures. Its release from the plasma membrane potentially reflects the activation of EMT-like processes in lung organoids, accompanied by the decrease of expressed *KRT8*. Further supporting these findings, we showed the elevated expression of *BMP4* and *MMP7*, critical regulators of EMT in pulmonary fibrosis (Molloy et al., 2008), in lung organoids co-cultured with ILD fibroblasts. *MMP7* has furthermore been highlighted for its function as a plasma biomarker in idiopathic pulmonary fibrosis (Adams et al., 2020; Bauer et al., 2017), in line with its detection in our study.

Co-culture of alveolar organoids with ILD fibroblasts also specifically changed the expression of genes involved in ECM biosynthesis including the increased transcription of *Col1A1, VIM* and *CDH2*, well-known players in fibrotic lung disease (Adams et al., 2020; Katzen & Beers, 2020) (Ng et al., 2022), as compared to organoid monocultures. Linking back to the indications of aberrant basal transdifferentiation in fibroblast co-culture discussed above (Kathiriya et al., 2022), the basaloid cells show increased ECM protein expression, next to their increase in *BMP4* and *ITGB6* expression, again successfully detected in our model.

We next were able to provide deeper insight into the relevant mediators of mesenchymal-to-epithelial crosstalk related to the observed changes in organoid phenotype by screening the supernatant of ILD or control fibroblasts using mass spectrometry (Strieter et al., 2007). Cytokines that belong to the C-X-C motif family dominated the protein signature in the secretome, demonstrating their increased abundance in ILD. These signaling molecules act on the *CXCR1* and *CXCR2* receptor, as well as regulate the expression of cytokines from the interleukin family, central to the pathogenesis of fibrotic and inflammatory lung diseases such as IPF and acute respiratory distress syndrome (Kortekaas et al., 2022; Mukaida, 2003; Ng et al., 2020; Strikoudis et al., 2019). The majority of the differentially abundant proteins are proinflammatory cytokines that primarily act on neutrophil, monocyte or lymphocyte recruitment (*CXCL1, CXCL3, CXCL5, CXCL8*). The proinflammatory response is complemented by the regulation of proteins that play a role in cellular senescence and activation of *TGF-ß* such as the Pregnancy-Specific Glycoprotein (PSG) family, *PSG 4, 5, 6* or in the induction of a hypercoagulable tissue state (*TFPI2*). Other important proteins such as *SFRP4* directly inhibit WNT signaling, thereby modulating cell growth and differentiation, particularly of AT2s to AT1s (Abdelwahab et al., 2019). *WNT* serum levels are discussed as biomarkers for lung fibrosis and EMT. *WNT* modulation and *TGF-ß* solubilization in particular could account for the change in organoid number and size and support our observations in gene expression levels that indicate epithelial transdifferentiation into aberrant basaloid cells or EMT in the presence of ILD fibroblasts.

*IL11*, centrally orchestrating the ILD protein profile as observed by pathway enrichment, was identified among the top 15 candidates in the ILD fibroblast secretome. *IL11* is known to be expressed in pro-inflammatory fibroblasts extracted from IPF lungs. The cytokine belongs to the *IL6* family and is induced by *TGF-ß* and other proinflammatory mediators (*IL1β, IL17, IL22*, reactive oxygen species (ROS)). It can either activate fibroblasts to differentiate into myofibroblasts in an autocrine fashion through ERK/SMAD canonical signaling, which results in pro-fibrotic protein expression (*COL1A1, ACTA2*), or it stimulates epithelial cells (paracrine loop) through activating *ERK* signaling cascades, thereby inducing cellular senescence, EMT, cellular dysfunction and impaired regeneration (Ng et al., 2020). Results from our co-culture model indicate both autocrine and paracrine signaling of *IL11* as we demonstrated indication of EMT, stem cell dysfunction as well as ECM production, i.e., upregulation of collagen expression in line with Ng et al., 2020 (Ng et al., 2020). Similar to the results obtained from ILD fibroblast co-culture in different seeding densities, *IL11* treatment of alveolar organoid monocultures resulted in a dose dependent increase in metabolic activity, elevated expression of mesenchymal markers and decreased AT2 stemness and identity. Dose-dependent, *IL11* even induced apoptotic cell death, in line with its role in senescence and stem cell function observed in alveolar organoids (Kortekaas et al., 2022). Similarly, *IL11* alone induced fibrotic changes in healthy alveolar organoids whereas knock-out of *IL11* expression in diseased organoids reversed organoid fibrosis in a model of the Hermansky-Pudlak syndrome-associated interstitial pneumonia, a disease with high similarity to IPF (Strikoudis et al., 2019). Supporting our co-culture findings in this regard, *IL11* exposure impacts on AT2 progenitor function, thereby likely suppressing the formation of mature AT2s as described by Kortekaas and colleagues (Kortekaas et al., 2022; Kortekaas et al., 2021).

Taken together, our results strongly support the central role of *IL11* signaling as the cytokine holds the potential to strongly influence the intricate crosstalk between the (then activated (myo)) fibroblasts and the injured epithelium, central in the progression of fibrosis in ILD. We successfully demonstrated the potential of our lung organoid co-culture model derived from hiPSCs and primary fibroblasts to display critical consequences of the malfunctional crosstalk such as aberrant dedifferentiation and basaloid-prone signatures (Adams et al., 2020; Habermann et al., 2020). In this context, *IL11* likely holds an important role in misguided alveolar function, differentiation and thereby regeneration, important functions as a potential therapeutic target to regain alveolar crosstalk homeostasis (Lin et al., 2022).

## Author Contributions

M.T. Kastlmeier, A. Hilgendorff and C. Voss conceived and planned the experiments. M.T. Kastlmeier and E. Gonzalez Rodriguez carried out the experiments. M.T. Kastlmeier, E. Gonzalez Rodriguez, P. Cabanis, E. M. Guenther contributed to sample preparation. A-C. König and S. Hauck performed the mass spectrometry analysis. M.T. Kastlmeier, C. Voss, T. Stoeger, A-C. König, S. Hauck and A. Hilgendorff contributed to the interpretation of the results. M.T. Kastlmeier and C. Voss. took the lead in writing the manuscript. All authors provided critical feedback and helped shape the research, analysis and manuscript.

## Acknowledgments

We gratefully acknowledge the provision of human biomaterial (primary human fibroblasts) and clinical data from the CPC-M bioArchive and its partners at the Asklepios Biobank Gauting, the LMU Hospital and the Ludwig-Maximilians-Universität München. We thank the patients and their families for their support. We wish to thank the Kotton Lab especially Prof. Darrell Kotton by providing the hiPSC cell line. We are grateful to David Kutschke (LHI, Helmholtz Center Munich) for technical assistance. We thank Dr. Kenji Schorpp and Dr. Kamyar Hadian (Research Unit Signaling and Translation, Molecular Targets and Therapeutics Center, Helmholtz Zentrum München, Germany) for their excellent support with the Operetta System. We thank the Research Unit Analytical Pathology (AAP, Helmholtz Center Munich) with Dr. Ulrike Buchholz and Dr. Annette Feuchtinger for their assistance during confocal imaging. We thank Benoite Champon and Dr. Minodora Brimpari for their help with the cell sorting at the MACSQuant Tyto Cell Sorter. We thank Zeynep Ertüz for creating the graphical abstract. Graphics were created with BioRender.com. The manuscript has been published as a preprint on Biorxiv under (Kastlmeier, Rodriguez, et al., 2022).

## Supplementary Material

### 1 Supplementary Figures and Tables

#### 1.1 Supplementary Tables

**Supplementary Table 1.** Antibodies and dilutions for immunofluorescence staining.

| Antibody | Manufacturer | Host Species | Isotype | Reactivity | Clonality | Dilution |
| --- | --- | --- | --- | --- | --- | --- |
| ALB | R&D Systems<br>Cat# MAB1455<br>Clone# 188835 | mouse | IgG2a | human, mouse, rat | monoclonal | 1:50 |
| α- SMA | Abcam<br>Cat# ab124964<br>Lot# GR303485-6 | rabbit | NA | human, mouse, rat | monoclonal | 1:200 |
| CPM | Fujifilm Wako<br>Chemicals<br>Cat# 014-27501 | mouse | IgG2b | human | monoclonal | 1:200 |
| NKX2.1 | Invitrogen<br>Cat# MA5-13961<br>Clone# 8G7G3/1 | mouse | IgG1 $\kappa$ | human, mouse, rat | monoclonal | 1:50 |
| Pro SP-C | Millipore<br>Cat # AB3786<br>Lot# 2464523 | rabbit | IgG | human, mouse, rat | polyclonal | 1:200 |
| Alexa Flour 488 | Invitrogen | anti-mouse | IgG1 |  |  | 1:500 |
| Alexa Flour 647 | Invitrogen | anti-rabbit |  |  |  | 1:500 |
| DAPI | Sigma |  |  |  |  | 1:1000 |

**Supplementary Table 2.**
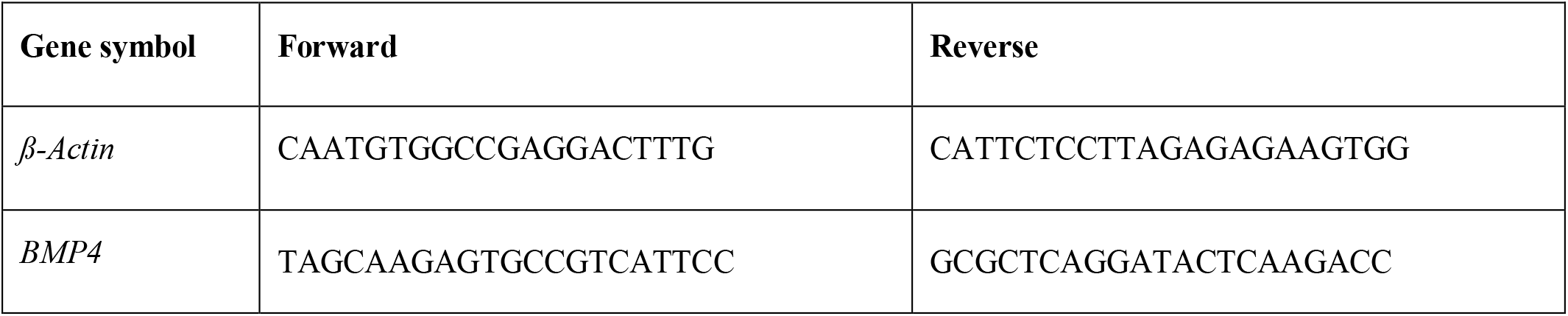
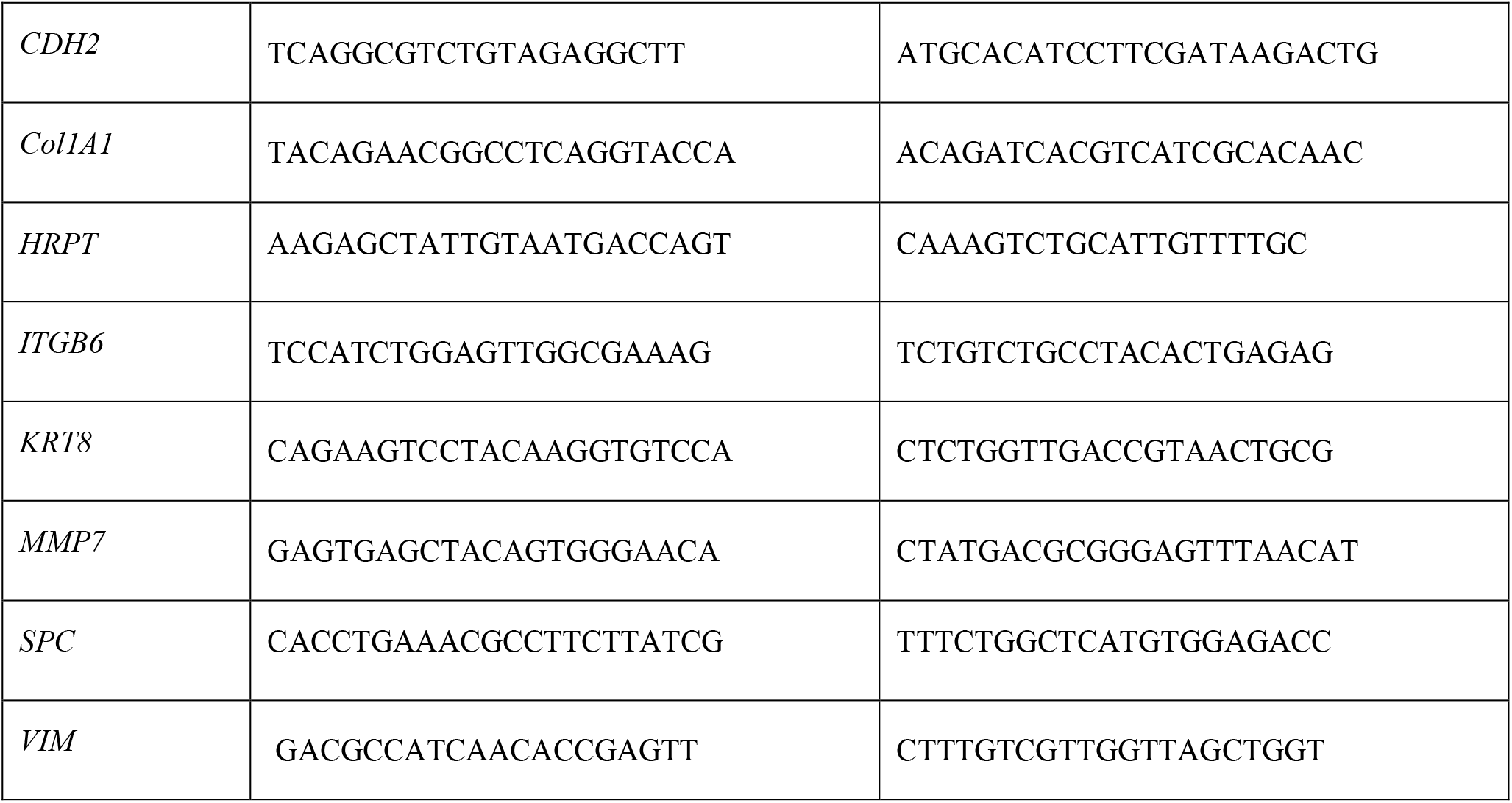
Primer sequences for qPCR.

**Supplementary Table 3.** List of upregulated proteins compared F_Control_ vs. F_ILD_ identified by mass spectrometry.

| Protein<br>FDR<br>confidence | Accession | Unique<br>peptides | Gene<br>symbol | Description | Abundance<br>ratio | Abundance<br>ratio P-value | Abundance<br>ratio adj.<br>P-value |
| --- | --- | --- | --- | --- | --- | --- | --- |
| High | Q68BL8 | 2 | OLFML2B | Olfactomedin-like protein 2B | 100,00 | 0,0000 | 0,0000 |
| High | Q9C0H2 | 3 | TTYH3 | Protein tweety homolog 3 | 100,00 | 0,0000 | 0,0000 |
| High | Q96FV2 | 2 | SCRN2 | Secernin-2 | 100,00 | 0,0000 | 0,0000 |
| High | Q9UHX1 | 3 | PUF60 | Poly(U)-binding-splicing factor PUF60 | 100,00 | 0,0000 | 0,0000 |
| High | Q16630 | 2 | CPSF6 | Cleavage and polyadenylation specificity factor subunit 6 | 100,00 | 0,0000 | 0,0000 |
| High | Q00888 | 8 | PSG4 | Pregnancy-specific beta-1-glycoprotein 4 | 8,81 | 0,0000 | 0,0000 |
| High | P17302 | 3 | GJA1 | Gap junction alpha-1 protein | 7,85 | 0,0000 | 0,0000 |
| High | P09341 | 2 | CXCL1 | Growth-regulated alpha protein | 7,06 | 0,0000 | 0,0000 |
| High | P19876 | 3 | CXCL3 | C-X-C motif chemokine 3 | 6,42 | 0,0000 | 0,0000 |
| High | Q14956 | 2 | GPNMB | Transmembrane glycoprotein NMB | 6,02 | 0,0000 | 0,0000 |
| High | P48307 | 11 | TFPI2 | Tissue factor pathway inhibitor 2 | 4,92 | 0,0000 | 0,0000 |
| High | Q14627 | 3 | IL13RA2 | Interleukin-13 receptor subunit alpha-2 | 4,46 | 0,0000 | 0,0000 |
| High | P20809 | 3 | IL11 | Interleukin-11 | 4,31 | 0,0000 | 0,0000 |
| High | Q6FHJ7 | 2 | SFRP4 | Secreted frizzled-related protein 4 | 4,05 | 0,0000 | 0,0001 |
| High | P10145 | 2 | CXCL8 | Interleukin-8 | 3,75 | 0,0000 | 0,0000 |

**Supplementary Table 4.** List of downregulated proteins compared F_Control_ vs. F_ILD_ identified by mass spectrometry.

| **Protein FDR confidence** | **Accession** | **Unique peptides** | **Gene symbol** | **Description** | **Abundance ratio** | **Abundance ratio P-value** | **Abundance ratio adj. P-value** |
| --- | --- | --- | --- | --- | --- | --- | --- |
| High | Q15198 | 5 | PDGFRL | Platelet-derived growth factor receptor-like protein | 0,49 | 0,0025 | 0,0325 |
| High | P10321 | 4 | HLA-C | HLA class I histocompatibility antigen, C alpha chain | 0,47 | 0,0012 | 0,0172 |
| High | P22352 | 8 | GPX3 | Glutathione peroxidase 3 | 0,43 | 0,0002 | 0,0034 |
| High | Q9NZP8 | 7 | C1RL | Complement C1r subcomponent-like protein | 0,42 | 0,0001 | 0,0024 |
| High | P24821 | 87 | TNC | Tenascin | 0,42 | 0,0043 | 0,0500 |
| High | Q9NR99 | 36 | MXRA5 | Matrix-remodeling-associated protein 5 | 0,42 | 0,0041 | 0,0488 |
| High | Q13822 | 29 | ENPP2 | Ectonucleotide pyrophosphatase/p hosphodiesterase family member 2 | 0,42 | 0,0041 | 0,0485 |
| High | Q76M96 | 25 | CCDC80 | Coiled-coil domain-containing protein 80 | 0,41 | 0,0039 | 0,0473 |
| High | O43854 | 20 | EDIL3 | EGF-like repeat and discoidin I-like domain-containing protein 3 | 0,41 | 0,0038 | 0,0457 |
| High | Q9NRN5 | 13 | OLFML3 | Olfactomedin-like protein 3 | 0,41 | 0,0038 | 0,0458 |
| High | P02461 | 61 | COL3A1 | Collagen alpha-1(III) chain | 0,41 | 0,0035 | 0,0434 |
| High | P62906 | 7 | RPL10A | 60S ribosomal protein L10a | 0,41 | 0,0031 | 0,0391 |
| High | P00746 | 15 | CFD | Complement factor D | 0,41 | 0,0031 | 0,0391 |
| High | O00339 | 17 | MATN2 | Matrilin-2 | 0,40 | 0,0002 | 0,0039 |
| High | O15232 | 11 | MATN3 | Matrilin-3 | 0,40 | 0,0001 | 0,0023 |

#### 1.2 Supplementary Figures

**Supplementary Figure 1.**
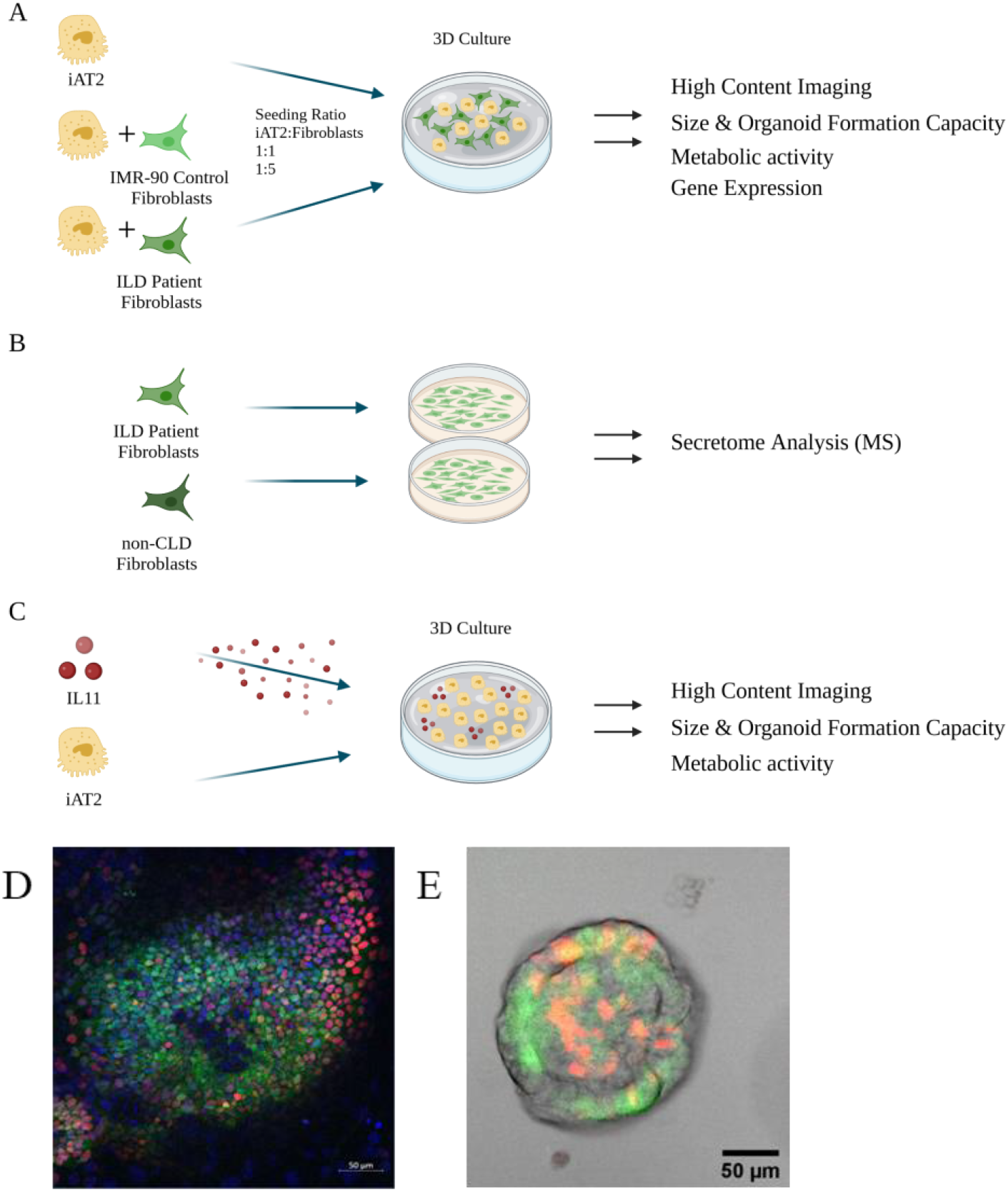
(A-C) Schematic workflow of the experiments. Created with BioRender.com. (D) Immunofluorescence of lung progenitors at day 14 of alveolar organoid differentiation (Albumin: red, NKX2.1: green, DAPI: cyan). (E) Alveolar organoids at day 44 of differentiation (BU3NGST cell line; *NKX2*.*1*^*GFP+*^, *SFTPC*^*tdTomato+*^).

**Supplementary Figure 2.**
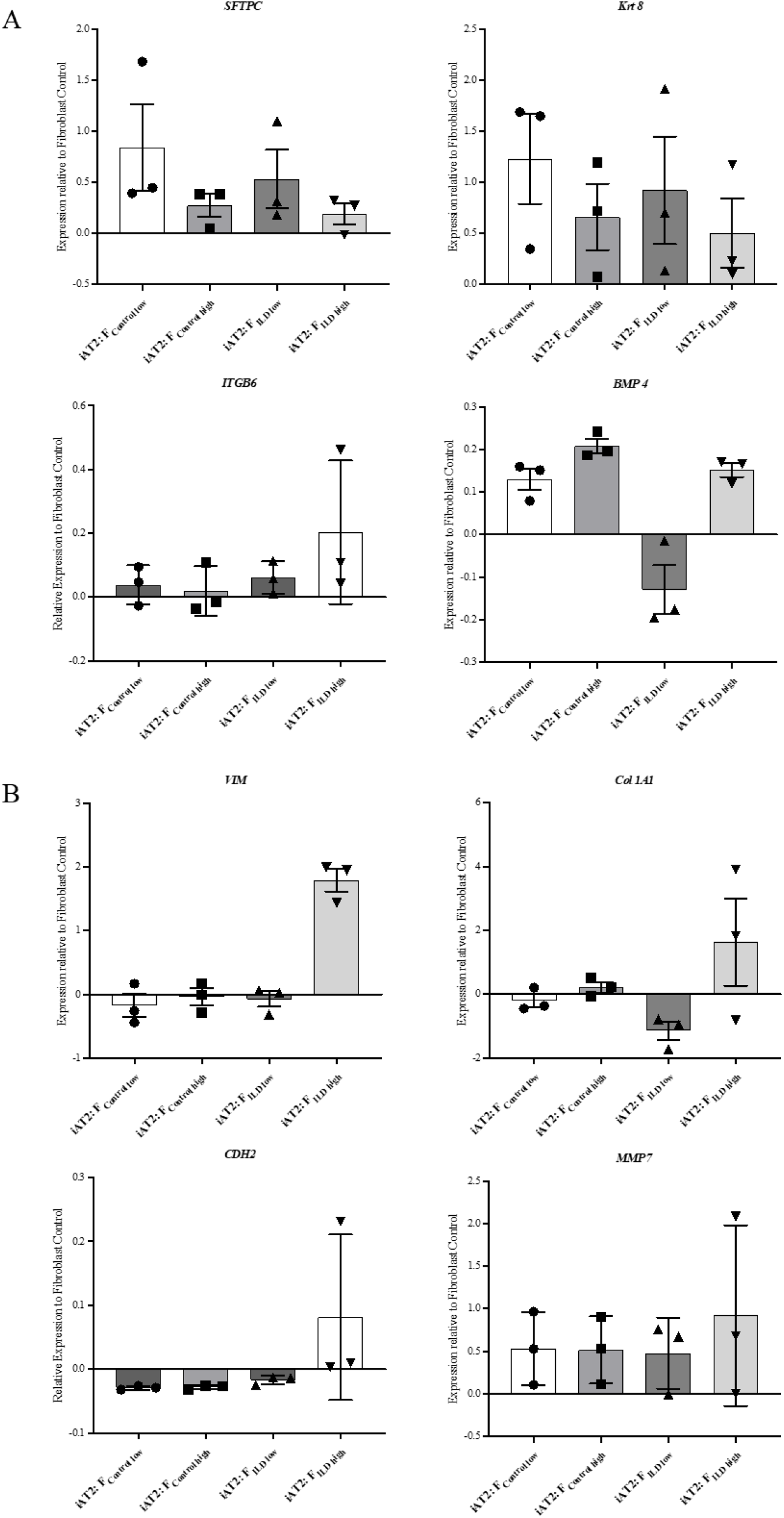
Comparison of relative gene expression of co-cultures with mono-cultured ILD or IMR90 control fibroblasts to understand potential epithelial origin of observed gene expression changes in co-cultures. Co-cultures were obtained by culturing iAT2s with either IMR-90 control fibroblasts or ILD fibroblasts in two seeding ratios (F_low_ and F_high_), N = 3. (A) Epithelial and stem cell markers and (B) genes associated with aberrant differentiation of epithelium.

